# Synthesizing and Patterning Tunable Multiscale Materials with Engineered Biofilms

**DOI:** 10.1101/002659

**Authors:** Allen Y. Chen, Urartu O. S. Seker, Michelle Y. Lu, Robert J. Citorik, Timothy K. Lu

## Abstract

A major challenge in materials science is to create self-assembling, functional, and environmentally responsive materials which can be patterned across multiple length scales. Natural biological systems, such as biofilms, shells, and skeletal tissues, implement dynamic regulatory programs to assemble complex multiscale materials comprised of living and non-living components^1–9^. Such systems can provide inspiration for the design of heterogeneous functional systems which integrate biotic and abiotic materials via hierarchical self-assembly. Here, we present a synthetic-biology platform for synthesizing and patterning self-assembled functional amyloid materials across multiple length scales with bacterial biofilms. We engineered *Escherichia coli* curli amyloid production under the tight control of synthetic regulatory circuits and interfaced amyloids with inorganic materials to create a biofilm-based electrical switch whose conductance can be selectively toggled by specific environmental signals. Furthermore, we externally tuned synthetic biofilms to build nanoscale amyloid biomaterials with different structure and composition through the controlled expression of their constituent subunits with artificial gene circuits. By using synthetic cell-cell communication, our engineered biofilms can also autonomously manufacture dynamic materials whose structure and composition change with time. In addition, we show that by combining subunit-level protein engineering, controlled genetic expression of self-assembling subunit proteins, and macroscale spatial gradients, synthetic biofilms can pattern protein biomaterials across multiple length scales. This work lays a foundation for synthesizing, patterning, and controlling composite materials with engineered biological systems. We envision that this approach can be expanded to other cellular and biomaterials contexts for the construction of self-organizing, environmentally responsive, and tunable multiscale composite materials with heterogeneous functionalities.

---

An overarching goal of synthetic biology is to re-design natural biological systems for useful applications. Synthetic cellular systems have been engineered to secrete protein materials such as silk^10^ and engage in intercellular communication to produce spatial patterns^11–14^. Natural multicellular assemblies such as biofilms, shells, and skeletal tissues have distinctive characteristics that would enable useful material production and patterning capabilities^1–9^. They have sophisticated mechanisms for detecting external signals and responding via remodelling, for implementing hierarchical patterning at different length scales, and for organizing inorganic compounds to create organic-inorganic composites. To harness the power of these features, we designed artificial gene circuits and engineered bacterial amyloid fibrils to achieve these multifunctional characteristics in the context of a synthetic biofilm system.

Our model system is curli, an extracellular amyloid protein material produced by *Escherichia coli* that forms fibrils based on the self-assembly of the major curli subunit CsgA^15^. Self-assembling proteins are structural molecules with rich chemical functionality which can be genetically encoded, not only making them amenable to engineering but also allowing their production to be controlled by synthetic regulatory elements in living cells. We implemented chemical-inducer-dependent synthetic transcriptional and translational control over the expression of CsgA, linking production of protein materials to environmental signals sensed by living cells. We used this synthetic circuit regulating CsgA expression to bring useful characteristics of natural multicellular assemblies to material production and patterning. By combining synthetic control of CsgA production with the ability of functionalized curli fibrils to interface with inorganic materials, we created artificial biofilms that act as electrical switches that are selectively toggled by a specific environmental signal. Moreover, through the use of inducer signals that have deliberately tuned temporal interval lengths and amplitudes, we achieved the production of materials with tunable structure and composition. Combining synthetic control of CsgA production with synthetic cell-cell communication allowed the generation of a dynamic material whose structure and composition changes autonomously with time. The introduction of protein engineering and macroscale spatial inducer gradients resulted in patterning across multiple length scales. In these ways, synthetic genetic regulatory elements enable the creation of engineered biological factories for environmentally responsive, tunable multiscale materials patterning.

To implement chemical inducer-dependent synthetic transcriptional and translational control over expression of CsgA, we knocked out the endogenous *csgA* gene from *E. coli* and introduced CsgA or histidine-tagged CsgA (CsgA-His) expression under tight regulation by an acyl-homoserine lactone (AHL) inducer-responsive or an anhydrotetracycline (aTc) inducer-responsive riboregulator^16^. The resulting cell strains were designated AHL-Receiver/CsgA, AHL-Receiver/CsgA-His, aTc-Receiver/CsgA, and aTc-Receiver/CsgA-His. We showed that curli fibrils were only produced in the presence of the proper inducer (Fig. 1a), and verified that insertion of heterologous histidine tags into CsgA did not interfere with curli fibril production (Supplementary Fig. 1).

**Figure 1.**
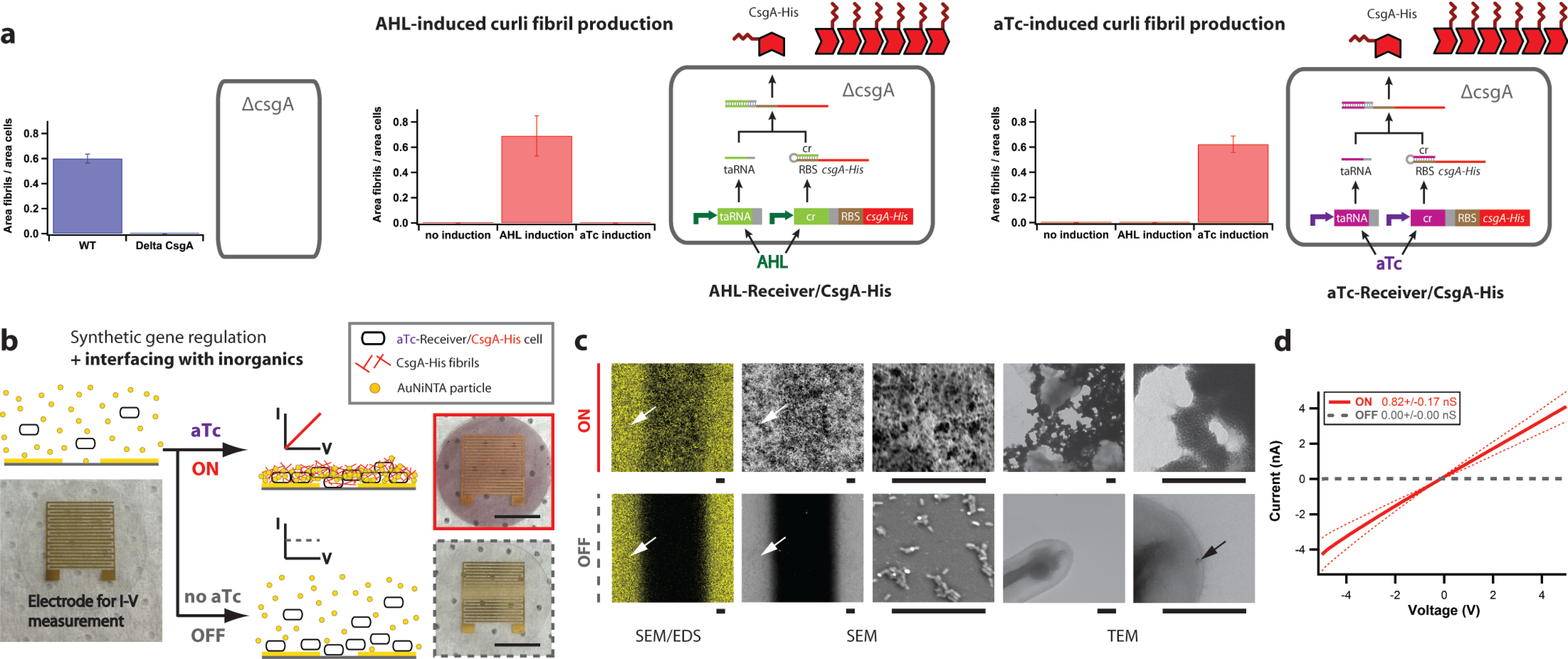
Environmentally switchable conductive biofilms. **a**, Engineered *E. coli* only produce curli fibrils when induced by the correct inducer. The curli knockout host strain used to create engineered inducer-responsive strains (MG1655 *PRO* Δ*csgA ompR234*) does not produce curli fibrils. The AHL-responsive engineered strain (AHL-Receiver/CsgA-His) only produces curli fibrils when induced by AHL. The aTc-responsive engineered strain (aTc-Receiver/CsgA-His) only produces curli fibrils when induced by aTc. Curli fibril production was quantified by taking the ratio of area of fibrils to area of cells in TEM images analysed by ImageJ. **b**, We created cells with synthetic genetic regulatory circuits that are regulated through external chemical inducer signals, such as anhydrotetracycline (aTc), to form amyloid fibrils via the production of His-tagged CsgA subunits (aTc-Receiver/CsgA-His cells). When combined with Ni-NTA gold particles (AuNiNTA particles), this implements an external signal-toggled conductive biofilm that acts as an electrical switch. When aTc inducer was added to aTc-Receiver/CsgA-His cells grown in the presence of AuNiNTA particles, it triggered the formation of conductive biofilms on electrodes. Scale bars are 5 mm. **c**, SEM/EDS elemental mapping of the aTc-induced ‘ON’ state for aTc-Receiver/CsgA-His biofilms showed that networks of gold in the biofilm connected electrodes (white arrows), SEM imaging showed that the biofilm bridged electrodes, and TEM imaging showed networks of aggregated gold particles. In contrast, SEM/EDS mapping of the ‘OFF’ state showed no gold networks, SEM imaging showed only scattered cells in the gap between electrodes, and TEM imaging showed only scattered and isolated gold particles (black arrow). Scale bars of scanning electron micrographs are 20μm and scale bars of transmission electron micrographs are 200 nm. **d**, The aTc-induced ‘ON’ state for aTc-Receiver/CsgA-His biofilms has a conductance of 0.82 ± 0.17 (s.e.m.) nS (solid red line), while the ‘OFF’ state has no measureable conductance (dashed grey line).

To create environmentally switchable conductive biofilms, we used the sensing functionality provided by the synthetic regulatory circuit in aTc-Receiver/CsgA-His cells, and the nickel nitrilotriacetic acid (Ni-NTA) binding functionality endowed by histidine tags to interface with Ni-NTA gold particles (AuNiNTA particles). The extracellular curli fibrils assembled from CsgA-His contribute structural integrity and surface-adherence functionality to the multicellular bacterial community, resulting in a thick adherent biofilm that forms on surfaces (Supplementary Fig. 2). Amyloid fibrils are advantageous as the backbone for engineered biofilms due to beneficial materials properties such as resistance to degradation and mechanical strength comparable to that of steel^17^.

We hypothesized that CsgA-His monomers secreted by aTc-Receiver/CsgA-His cells would self-assemble into extracellular amyloid fibrils that could organize AuNiNTA particles into chains to form a conductive biofilm network. Engineered biofilms were grown on interdigitated electrodes deposited on Thermanox coverslips, with aTc-Receiver/CsgA-His cells cultured in the presence of AuNiNTA particles and either in the presence or absence of aTc inducer (Fig. 1b). Scanning electron microscopy (SEM), scanning electron microscopy/energy dispersive X-ray spectroscopy (SEM/EDS), and transmission electron microscopy (TEM) were performed to characterize biofilm samples (Fig. 1c). SEM imaging of biofilms formed in the presence of inducer showed thick biofilm that spanned electrodes, and SEM/EDS elemental mapping showed networks of gold in the biofilm that connected electrodes. In contrast, SEM imaging of biofilms formed in the absence of chemical induction showed only scattered cells in the gaps between electrodes, and SEM/EDS showed no gold networks. TEM imaging revealed that the thick biofilms triggered by chemical induction organized gold particles into dense networks, while biofilms formed in the absence of chemical induction showed only scattered, isolated gold particles (Fig. 1c). Biofilms formed in the presence of inducer had 0.82 ± 0.17 (s.e.m.) nanosiemens conductance, whereas biofilms formed in the absence of inducer had no measureable conductance (Fig. 1d). Biofilms formed with aTc-Receiver/CsgA-His cells induced by aTc, but grown in the absence of AuNiNTA particles, had conductance two orders of magnitude lower than those formed in presence of AuNiNTA particles (Supplementary Fig. 3). Biofilms formed from AHL-Receiver/CsgA-His cells grown in the presence of AuNiNTA particles and aTc had no measureable conductance (Supplementary Fig. 3).

We next engineered biofilms to produce two-component protein fibrils whose nanoscale structure and composition are controlled via synthetic genetic regulatory elements (Fig. 2). These regulatory elements translate both the temporal interval length and the amplitude of external chemical inducer signals into different structure and composition of fibrils. Our two-component fibrils are formed from tunable combinations of CsgA and CsgA-His monomers. This approach could be generalized to include multiple CsgA variants with different properties and functionalities endowed by fusing various functional peptides. For CsgA monomer secretion, we used AHL-Receiver/CsgA and for CsgA-His monomer secretion, we used aTc-Receiver/CsgA-His (Fig. 2). To show that the structure of our two-component fibril is responsive to external signals, we mixed equal numbers of AHL-Receiver/CsgA and aTc-Receiver/CsgA-His, cultured cells in glycerol-supplemented M63 media, and induced this mixed-cell population first with AHL, followed by aTc (Fig. 2a). This produced block “co-fibrils” (in analogy to block co-polymers) consisting of blocks of CsgA (unlabeled fibril segments) and blocks of CsgA-His (fibril segments labeled by AuNiNTA particles).

**Figure 2.**
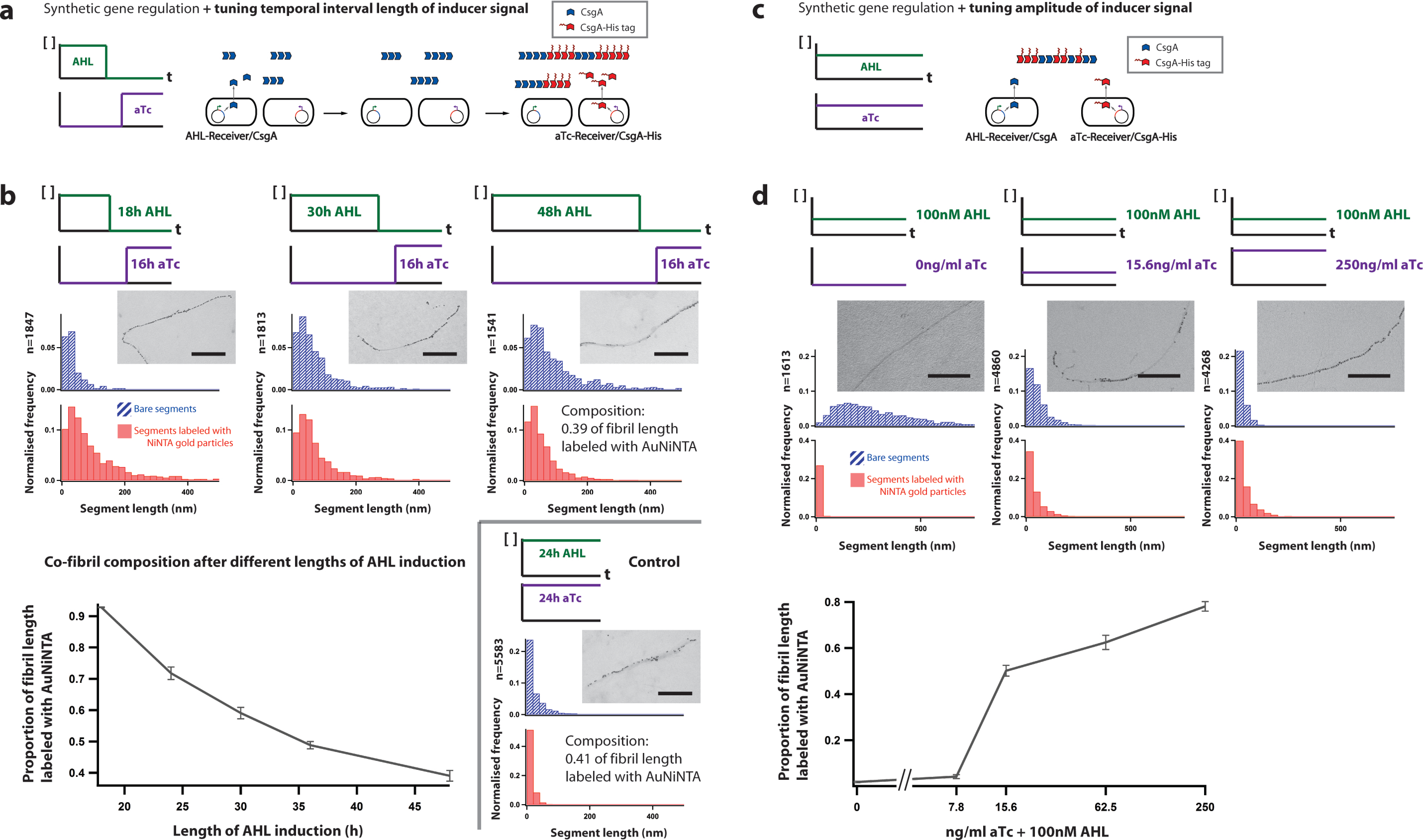
Conversion of timing and amplitude of chemical-inducer signals into material structure and composition. **a**, Synthetic genetic regulatory circuits that couple curli subunit secretion to external chemical inducer signals, when combined with inducer signals with deliberately tuned temporal interval lengths, allow production of amyloid fibrils with tunable structure and composition. **b**, With synthetic gene circuits, cells can translate the temporal interval length of input signals into nanoscale structure and composition of block co-fibrils. We first used AHL to induce secretion of CsgA from AHL-Receiver/CsgA, then used aTc to induce secretion of CsgA-His from aTc-Receiver/CsgA-His. We tuned the length distribution of the CsgA and CsgA-His blocks (histograms showing the length distributions of unlabeled and AuNiNTA labeled segments), as well as the relative proportions of CsgA and CsgA-His (plot of proportion of fibril length labeled by AuNiNTA, solid grey line), by changing the relative length of AHL versus aTc induction times. Hashed blue bars indicate bare segments of amyloid fibrils that were unlabeled by AuNiNTA, while solid red bars indicate amyloid fibril segments labeled with AuNiNTA particles. Compared with co-fibril variants with similar ratios of CsgA:CsgA-His (Control), the segments in block co-fibrils were longer than those in co-fibrils assembled when CsgA and CsgA-His were secreted simultaneously with no temporal separation. Scale bars are 200 nm. **c**, Synthetic genetic regulatory circuits that couple curli subunit secretion to external chemical inducer signals, when combined with inducer signals with deliberately tuned amplitudes, allow production of amyloid fibrils with tunable structure and composition. **d**, With synthetic gene circuits, cells can translate the amplitude of input signals into nanoscale structure and composition of block co-fibrils. AHL induces secretion of CsgA from AHL-Receiver/CsgA, while at the same time aTc induces secretion of CsgA-His from aTc-Receiver/CsgA-His. We tuned the length distributions of the CsgA and CsgA-His blocks, as well as the relative proportions of CsgA and CsgA-His, by changing the relative concentration of AHL and aTc inducers. Hashed blue bars indicate bare segments of amyloid fibrils that were unlabeled, while solid red bars indicate amyloid fibril segments labeled with AuNiNTA particles. Solid grey line indicates proportion of fibril length labeled by AuNiNTA particles. Scale bars are 200 nm.

We could tune the length distribution of the CsgA and CsgA-His blocks, as well as the relative proportions of CsgA and CsgA-His, by changing the relative lengths of AHL versus aTc induction times. As the induction time by AHL increased, fibril segments that were not labeled by AuNiNTA particles increased in length, indicating longer CsgA blocks (Fig. 2b). At the same time, the proportion of fibril segment length labeled with AuNiNTA particles decreased, indicating a higher relative proportion of CsgA in the fibrils. The segments in block co-fibrils achieved with temporal separation in expression of CsgA and CsgA-His were longer than those in co-fibrils assembled when CsgA and CsgA-His were secreted simultaneously with no temporal separation, even though the overall ratios of CsgA:CsgA-His were similar (Fig. 2b). This data demonstrates that with synthetic gene circuits, cells can translate the temporal interval length of input signals into nanoscale structure and composition of materials.

We could also tune the length distributions of the two types of blocks, as well as the relative proportions of the two subunits, by culturing cells in glucose-supplemented M63 media and inducing simultaneously with different combinations of AHL and aTc concentration (Fig. 2c). With AHL-only induction, fibrils were almost uniformly unlabeled, and with increasing aTc concentration, the population as well as lengths of unlabeled fibril segments decreased while those of labeled fibril segments increased (Fig. 2d). With aTc-only induction, fibrils were almost uniformly labeled by AuNiNTA particles, and with increasing AHL concentration, the population as well as lengths of unlabeled segments increased (Supplementary Fig. 5). Thus, cells equipped with synthetic circuits can translate the amplitude of input signals (in the form of inducer concentration) into nanoscale structure and composition of materials.

We additionally implemented synthetic cell-cell communication circuits to create cells that autonomously produce a dynamic material whose structure and composition changes with time (Fig. 3). We engineered an *E. coli* strain that constitutively produces AHL at a low basal level and inducibly produces CsgA in the presence of aTc (AHL-Sender+aTc-Receiver/CsgA). We used this strain to communicate with another strain, AHL-Receiver/CsgA-His. To show that cell-cell communication allows for production of a dynamic material, we combined AHL-Sender+aTc-Receiver/CsgA cells and AHL-Receiver/CsgA-His cells in varying ratios and induced this mixed-cell population with aTc (Fig. 3a). Induction by aTc resulted in CsgA secretion. Over time, AHL accumulation led to increasing secretion of CsgA-His and thus generated an increased population as well as increased length of CsgA-His blocks, and a higher relative proportion of CsgA-His in material composition (Fig. 3b). The rate at which the material composition changed could be tuned by the initial seeding ratio of AHL-Sender+aTc-Receiver/CsgA cells to AHL-Receiver/CsgA-His cells. When only AHL-Sender+aTc-Receiver/CsgA cells were present, the resulting fibrils were almost uniformly unlabeled; when only AHL-Receiver/CsgA-His cells were present, no fibrils were formed (Fig. 3b).

**Figure 3.**
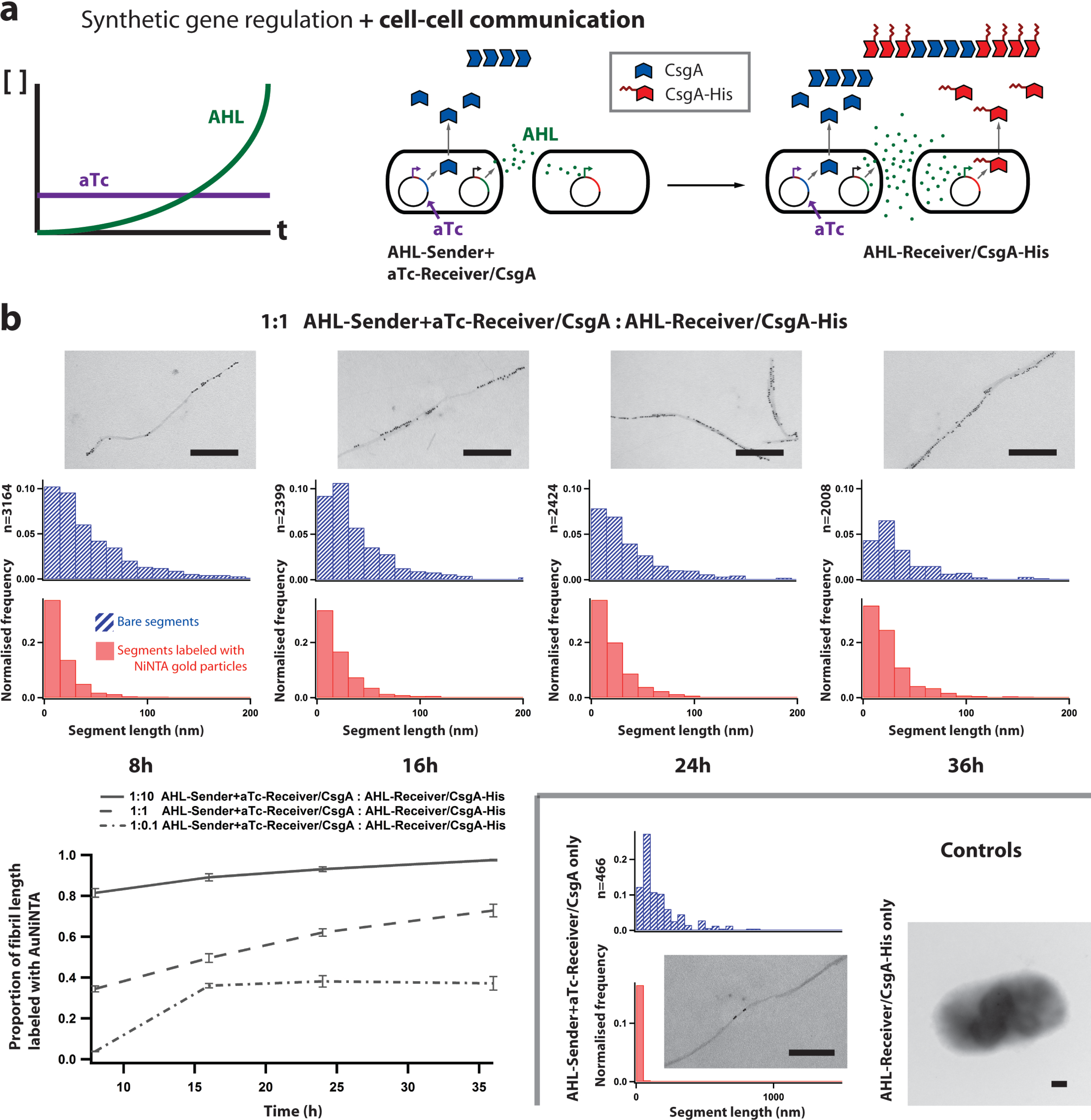
Synthetic cell-cell communication for dynamic and autonomous material production and patterning. **a**, Synthetic genetic regulatory circuits that couple curli subunit secretion to external chemical inducer signals, when combined with synthetic cell-cell communication, allow for the production of a dynamic material whose structure and composition change autonomously with time. AHL-Sender+aTc-Receiver/CsgA secretes both CsgA and AHL. As AHL signal accumulates, AHL-Receiver/CsgA-His secretes increasing levels of CsgA-His. **b**, Using the autonomous cell-cell communication system, the concentration of AHL increases over time, and the population as well as length of CsgA-His blocks increases with time (histogram of length distribution of unlabeled and AuNiNTA labeled segments). Hashed blue bars indicate bare segments of amyloid fibrils that were unlabeled, while solid red bars indicate amyloid fibril segments labeled with AuNiNTA particles. The proportion of CsgA-His increases with time (plot of proportion of fibril length labeled by AuNiNTA, grey lines) and could be tuned by the ratio of the seeding density of AHL-Sender+aTc-Receiver/CsgA cells to AHL-Receiver/CsgA-His cells. When only AHL-Sender+aTc-Receiver/CsgA cells were present, resulting fibrils were almost uniformly unlabeled (Controls); when only AHL-Receiver/CsgA-His cells were present, no curli fibrils were formed (Controls). Scale bars are 200 nm.

In addition to external or autonomous control of materials synthesis at the nanoscale, engineered biofilms can enable spatial control over multiple length scales. As described above, genetic regulation of subunit expression allows patterning of self-assembling fibrils from tens of nanometres to micrometres. Spatial control at the macroscale can be achieved via spatially varying input inducer concentrations. These two methods of control can be combined to create materials patterned at multiple length scales (Fig. 4).

**Figure 4.**
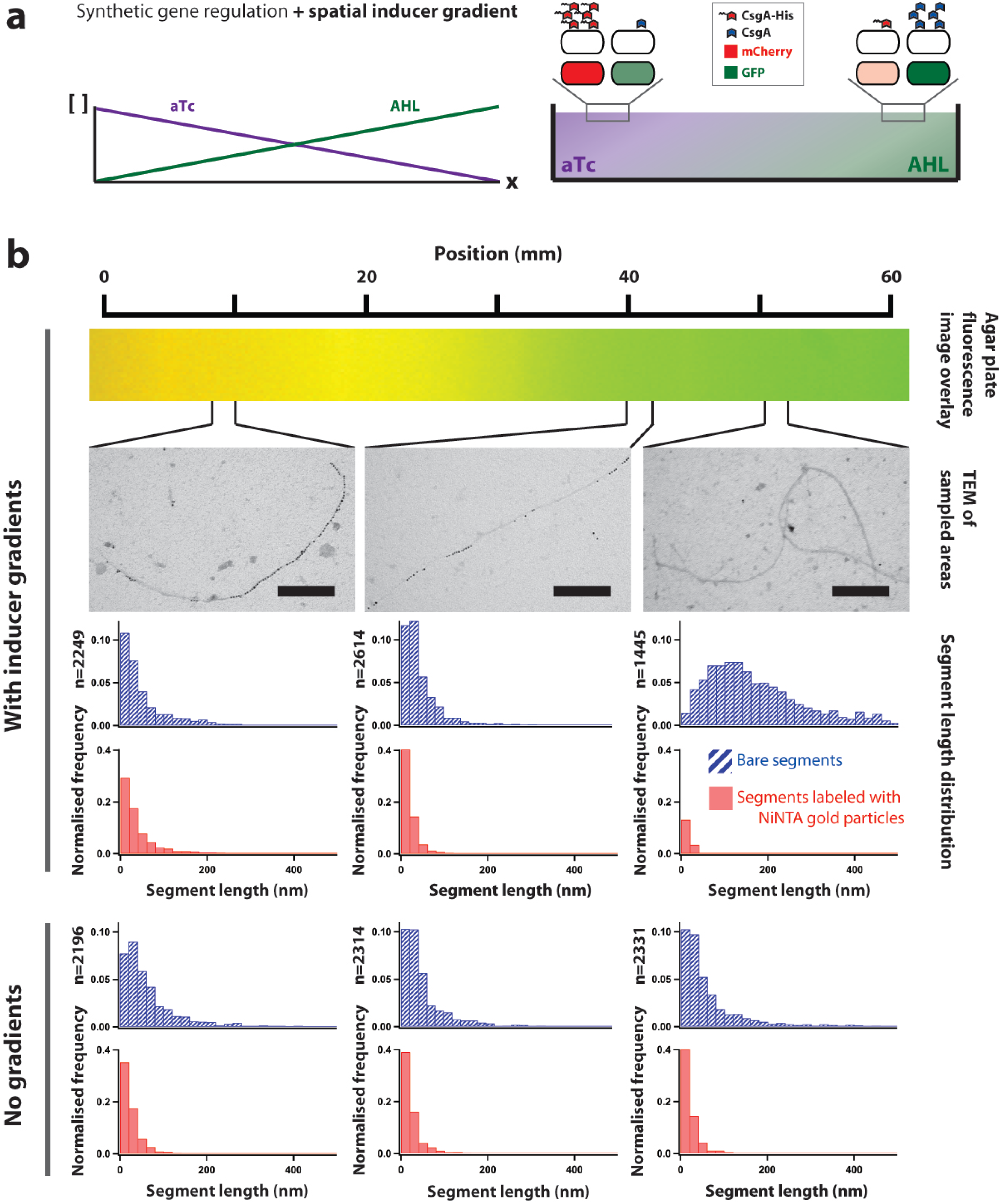
Multiscale patterning with engineered biofilms via synthetic gene regulation and inducer gradients. **a**, Synthetic regulatory circuits that couple curli subunit secretion to external chemical inducer signals, when combined with a spatial inducer gradient, enable spatial patterning across multiple length scales. We used an agar plate with opposing linear concentration gradients of AHL and aTc at the surface to achieve spatial control at the macroscale. This macroscale control was combined with synthetic gene regulation of nanoscale patterning to achieve multiscale patterning. Embedded in top agar were equal numbers of AHL-Receiver/CsgA, aTc-Receiver/CsgA-His, AHL-Receiver/GFP, and aTc-Receiver/mCherry cells. **b**, By combining synthetic gene regulation with spatial inducer gradients, we created a change in the nanoscale structure of fibrils across a distance of millimetres. This patterning at both the nanoscale and the macroscale was shown by changes in population and segment length of unlabeled and AuNiNTA labeled fibril segments at different locations across the agar plate. Inducer concentration gradients were demonstrated by overlaid GFP and mCherry fluorescence images of embedded reporter cells. Hashed blue bars indicate bare segments of amyloid fibrils that were unlabeled with AuNiNTA, while solid red bars indicate amyloid fibril segments labeled with AuNiNTA particles. Scale bars are 200 nm.

We created agar plates with opposing linear concentration gradients of AHL and aTc at the surface, and overlaid bacterial populations consisting of equal numbers of four cell strains on the agar: AHL-Receiver/CsgA, aTc-Receiver/CsgA-His, AHL-Receiver/GFP, and aTc-Receiver/mCherry. The AHL-Receiver/GFP and aTc-Receiver/mCherry cells enabled visualization of inducer concentration gradients (Supplementary Fig. 6 and Fig. 4b). AHL-Receiver/CsgA and aTc-Reciever/CsgA-His cells secreted different levels of CsgA and CsgA-His, depending on their positions on the concentration gradient, to generate a spatial gradient of changing fibril structure (Fig. 4a). This multiscale material was patterned at the nanoscale as block co-fibrils consisting of CsgA and CsgA-His and at the millimetre scale as position-dependent fibril structure (Fig. 4b). Agar plates without inducer concentration gradients did not generate fibril structures that varied along the agar plate (Fig. 4b).

Protein engineering provides another means of controlling the structure of cell-produced biomaterials at the nanoscale. We hypothesized that fusing together tandem repeats of CsgA would increase the distance between equivalent positions on adjacent monomers, such as the C-terminus, where functional domains can be attached. Concatenating eight tandem repeats of CsgA and adding a histidine tag to the C-terminus (8XCsgA-His) resulted in fibrils that were labeled by a syncopated pattern of AuNiNTA particles, with clusters of particles separated by 33.3 ± 27.1 (s.e.m.) nm (Fig. 5a). Using this finding, we demonstrated a second example of multiscale assembly. Specifically, we combined equal numbers of AHL-Receiver/8XCsgA-His and aTc-Receiver/CsgA-His cells and induced this mixed-cell population first with AHL, followed by aTc (Fig. 5b). This produced block co-fibrils consisting of segments of 8XCsgA-His and segments of CsgA-His patterned across the nanometre to micrometre scale (Fig. 5b).

**Figure 5.**
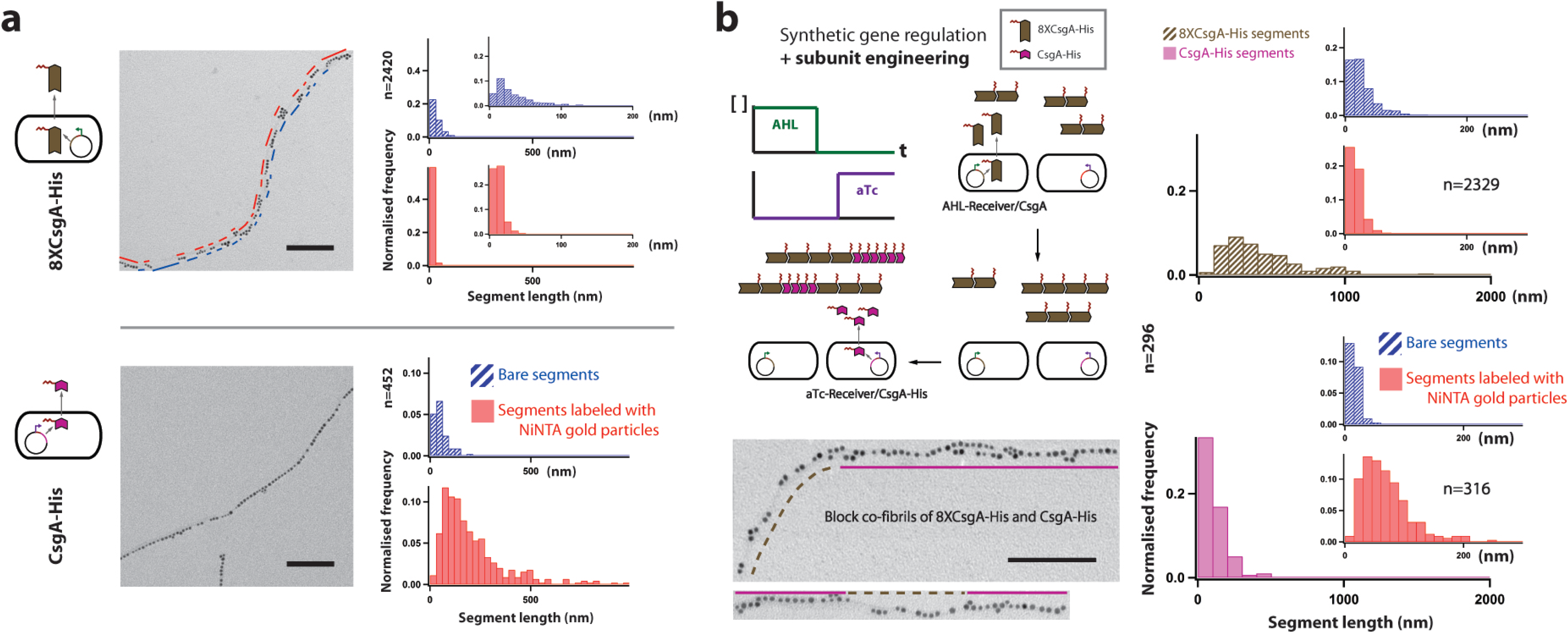
Multiscale patterning with engineered biofilms via synthetic gene regulation and subunit engineering. **a**, We achieved patterning at the nanoscale by protein engineering of curli subunits. Concatenating eight tandem repeats of CsgA and adding one histidine tag to the C-terminus (8XCsgA-His) resulted in fibrils that were labeled by a syncopated pattern of AuNiNTA particles, with clusters of particles separated by 33.3 ± 27.1 (s.e.m.) nm. Hashed blue bars indicate bare segments of amyloid fibrils that were unlabeled with AuNiNTA, while solid red bars indicate amyloid fibril segments labeled with AuNiNTA particles. Insets show smaller-binned histograms of the length distribution of unlabeled and labeled 8XCsgA-His fibril segments. Scale bars are 100 nm. **b**, Synthetic genetic regulatory circuits that couple curli subunit secretion to external chemical inducer signals, when combined with curli subunit engineering, enable spatial patterning across multiple length scales (nanometres to micrometres). We used AHL to induce production of 8XCsgA-His from AHL-Receiver/8XCsgA-His, then used aTc to induce production of CsgA-His from aTc-Receiver/CsgA-His. In the TEM images, dashed brown lines refer to syncopated 8XCsgA-His segments while the solid amethyst lines indicate CsgA-His segments. The length distributions of 8XCsgA-His segments and CsgA-His segments are shown in histograms with hashed brown bars and solid amethyst bars, respectively. Inset histograms show the length distributions of unlabeled segments (hashed blue bars) and segments labeled with AuNiNTA particles (solid red bars) for both the 8XCsgA-His segments and the CsgA-His segments. Scale bars are 100 nm.

We have demonstrated an engineered biofilm platform that uses artificial gene circuits to synthesize and pattern self-assembling materials with tunable functionality, structure, and composition. Since cells are capable of complex signal integration using cellular computation networks^18,19^, they can be used to assemble materials in response to environmental signals for a variety of applications. They can be “smart” factories that sense external conditions to produce the appropriate functional material at the right time, or programmable foundries that can receive instructions to generate materials with the desired function, structure, and composition. As cells are able to send and receive signalling molecules, they can be the basis for materials production platforms consisting of different types of cells that perform specialized functions and at the same time coordinate through cell-cell communication.

Living cells are natural platforms with which to achieve multiscale patterned materials because biology is organized in a hierarchical manner, from macromolecules (e.g. proteins, nucleic acids, carbohydrates, lipids) to macromolecular assemblies (synthetic variants of which are used as nanomaterials^20–23^) to organelles to cells and to tissues. In fact, natural biological materials such as bone are hierarchically organized to fulfil varied functional requirements^1,9^. The strategy described in this work uses synthetic gene circuits within engineered biofilms to achieve multiscale patterning of materials. This approach can be expanded to other cellular and biomaterials contexts for applications ranging from biointegrated electronic and optical devices^24–26^ to tissue engineering scaffolds^27^. For example, our demonstration of a gradient material patterned at the nanoscale and the millimetre scale could represent a fabrication strategy for functionally graded materials that complements top-down approaches for making such materials, such as 3D printing^28^. Mammalian cells capable of tunable, environmentally responsive synthesis of multiscale patterned materials may be used to mimic the dynamic microenvironment of the *in vivo* extracellular matrix, paving the way to engineered matrices that recapitulate tissue and organ function over time. Thus, the strategy of using the hierarchical organization of biology for multiscale patterning complements other top-down and bottom-up materials synthesis strategies that require directed human intervention^29^.

Beyond being a convenient model system with which to explore the applications of living systems to materials science, protein materials are of practical interest because they constitute a major class of biomaterials^30^, with programmable structure^23^ and diverse functionalities such as responsiveness to physicochemical stimuli^31^, the ability to interact with living systems^32^, and the ability to organize diverse abiotic materials for expanded functionality^33–36^. However, most existing examples of protein biomaterials are assembled *in vitro* from chemically synthesized peptides or purified subunits and do not take full advantage of the fact that the materials’ constituent subunits can be made by living cells. By integrating synthetic gene networks in engineered cells with extracellular protein biomaterials, living materials that possess specific environmental responsiveness, tunable functionality, multiscale patterning, and even the ability to self-heal and remodel could be realized. In such materials, there is a division of labour between cells (providing functionality of living systems)^37^, extracellular protein material (providing spatial patterning and structural integrity), and interfaced abiotic materials (providing functionality of non-living systems). Ultimately, we envision that engineering artificial cellular consortia, such as biofilms, to synthesize and organize heterogeneous functional materials will enable the realization of smart composite materials that combine the properties of living and non-living systems.

## Methods

### Plasmid construction

The plasmids used in this study were constructed with standard molecular cloning techniques^38^ using New England BioLabs (NEB) restriction endonucleases, T4 DNA Ligase, and Phusion PCR kits. A Bio-Rad S1000 Thermal Cycler with Dual 48/48 Fast Reaction Modules (Bio-Rad) was used to perform PCRs, ligations, and restriction digests. Gel extractions were carried out with QIAquick Gel Extraction Kits (Qiagen). Custom oligonucleotide primers were obtained from Integrated DNA Technologies (Coralville, IA).

All ligations for plasmid construction were transformed into *E. coli* strain DH5αPRO with standard protocols^38^. Isolated colonies were inoculated into Luria-Bertani (LB)-Miller medium (Fisher) using appropriate antibiotics at the following concentrations: carbenicillin at 50 μg/ml, chloramphenicol at 25 μg/ml. DNA was extracted with Qiagen QIAprep Spin Miniprep Kits. Plasmid construct sequences were confirmed by restriction digest and sequencing was performed by Genewiz (Cambridge, MA).

The parts that constitute the plasmids used in this work are described in Supplementary Table 1, and plasmids are described in Supplementary Table 2. To create constructs for expression of output genes under tight regulation by an aTc-inducible riboregulator, pZE-AmpR-rr12-pL(tetO)-*gfp* was used as a starting point^16^. The AmpR cassette of pZE-AmpR-rr12-pL(tetO)-*gfp* was swapped for the CmR cassette from pZS-CmR-pL(lacO)-*gfp* by using SacI-HF/AatII digestion and T4 ligation to create pZE-CmR-rr12-pL(tetO)-*gfp*. Then, the ColE1 origin of pZE-CmR-rr12-pL(tetO)-*gfp* was swapped for the p15A origin from pZA-AmpR-pL(tetO)-*gfp* by using SacI-HF/AvrII excision and T4 ligation to create pZA-CmR-rr12-pL(tetO)-*gfp*. The *gfp* was excised using KpnI/MluI digest to create the vector pZA-CmR-rr12-pL(tetO)-. The genes *csgA*, *csgA-His*, and *mCherry* with KpnI and MluI sticky ends were generated by PCR and KpnI/MluI digest; these fragments were ligated with pZA-CmR-rr12-pL(tetO)-to create pZA-CmR-rr12-pL(tetO)-*csgA*, pZA-CmR-rr12-pL(tetO)-*csgA-His*, and pZA-CmR-rr12-pL(tetO)-*mCherry* plasmids.

To create constructs for expression of output genes under tight regulation by an AHL-inducible riboregulator, pZE-KanR-rr12y-pLuxR-*gfp* was used as a starting point^16^. The KanR cassette of pZE-KanR-rr12y-pLuxR-*gfp* was swapped for the CmR cassette from pZS-CmR-pL(lacO)-*gfp* by using SacI-HF/AatII digestion and T4 ligation to create pZE-CmR-rr12y-pLuxR-*gfp*. Then the ColE1 origin of pZE-CmR-rr12y-pLuxR-*gfp* was swapped for the p15A origin from pZA-AmpR-pL(tetO)-*gfp* by using SacI-HF/AvrII excision and T4 ligation to create pZA-CmR-rr12y-pLuxR-*gfp*. The *gfp* was excised using KpnI/MluI digest to create the vector pZA-CmR-rr12y-pLuxR-. A codon-optimized gene encoding a CsgA variant consisting of eight tandem repeats of CsgA with a histidine tag at the C-terminus (*8XcsgA-His*), and flanked by KpnI and MluI sites, was designed and custom synthesized by GenScript (Piscataway, NJ). The gene *8XcsgA-His* with sticky ends was generated by KpnI/MluI digest and ligated with pZA-CmR-rr12y-pLuxR-to create pZA-CmR-rr12y-pLuxR-*8XcsgA-His*. The genes *csgA* and *csgA-His*, also with KpnI and MluI sticky ends, were ligated with the vector to create pZA-CmR-rr12y-pLuxR-*csgA* and pZA-CmR-rr12y-pLuxR-*csgA-His*.

An innocuous plasmid with an AmpR resistance marker was also created for use in a co-culture experiment. It has a low-copy origin and an innocuous output gene (*lacZ-alpha*) under tight repression by a riboregulator to ensure minimal effect on the host cell apart from conferring ampicillin resistance. The plasmid pZE-KanR-rr10-pL(lacO)-*mCherry* was used as a starting point. The KanR cassette of pZE-KanR-rr10-pL(lacO)-*mCherry* was swapped for the AmpR cassette from pZA-AmpR-pL(tetO)-*gfp* by using SacI-HF/AatII digestion and T4 ligation to create pZE-AmpR-rr10-pL(lacO)-*mCherry*. Then the ColE1 origin of pZE-AmpR-rr10-pL(lacO)-*mCherry* was swapped for the pSC101 origin from pZS-CmR-pL(lacO)-*gfp* by using SacI-HF/AvrII excision and T4 ligation to create pZS-AmpR-rr10-pL(lacO)-*mCherry*. The *mCherry* gene of pZSAmpR-rr10-pL(lacO)-*mCherry* was swapped for the *lacZ-alpha* gene from pZE-AmpR-pLuxR-*lacZ-alpha* using KpnI/MluI excision and T4 ligation to create pZS-AmpR-rr10-pL(lacO)-*lacZ-alpha*.

### Strain construction

To create the *E. coli* MG1655 *PRO* Δ*csgA ompR234* strain used in this study, we sequentially generated *E. coli* MG1655 *PRO* Δ*csgA::aph*, *E. coli* MG1655 *PRO* Δ*csgA*, and *E. coli* MG1655 *PRO* Δ*csgA ompR234*. The PRO cassette expresses *lacI* and *tetR* at high constitutive levels^39^.

*E. coli* MG1655 *PRO* Δ*csgA::aph* was generated via P1 transduction of the Δ*csgA::aph* allele from donor strain JW1025-1 of the Keio collection^40^ into recipient *E. coli* MG1655 *PRO*^16^, as previously described^38^. To prepare P1 lysate, 100μl overnight culture of donor strain was incubated with 100μl P1 phage in 10 ml LB+7 mM CaCl_2_+12 mM MgSO_4_ for 20 min without shaking at 37C, then for 2 h with shaking at 37C. To produce more P1 lysate, 300μl more of donor strain was added, the culture incubated for 20 min without shaking at 37C, then for 2 h with shaking at 37C. The culture was transferred to a 50 ml Falcon tube, 2 ml of chloroform was added, and the mixture was vortexed and centrifuged for 15 min at 3000 rpm. The supernatant, containing the P1 lysate, was transferred to Eppendorf tubes and centrifuged for 5 min at 14000 rpm to pellet remaining debris. For transduction, the P1 lysate was diluted 60-fold into 1 ml LB+7 mM CaCl_2_+12 mM MgSO_4_ in an Eppendorf tube, 100μl overnight culture of recipient strain was added, and the tube was incubated for 1 h with shaking at 37C. This culture was then plated onto LB kanamycin agar with 25 mM sodium citrate to select for recipients successfully transduced with the Δ*csgA::aph* allele. Transformants were restreaked onto LB kanamycin agar for re-isolation.

*E. coli* MG1655 *PRO* Δ*csgA* was generated by removing the kanamycin resistance cassette as previously described^41^ (in order to free this antibiotic selection marker for subsequent usage). Briefly, MG1655 *PRO* Δ*csgA::aph* was transformed with pCP20 carrying the gene for FLP recombinase and selected on LB ampicillin agar at 30°C. Transformants were restreaked onto non-selective LB and grown overnight at 42°C to cure the temperature-sensitive pCP20. Isolates were patched onto LB agar containing either kanamycin or ampicillin to confirm the loss of the FRT-flanked kanamycin cassette and pCP20, respectively.

Finally, *E. coli* MG1655 *PRO* Δ*csgA ompR234* was generated via P1 transduction of the *ompR234* allele (linked to a kanamycin resistance marker) from donor *E. coli* MG1655 *ompR234*^42^ into recipient *E. coli* MG1655 *PRO* Δ*csgA* with the same protocol as above. The final strain was verified via PCR amplicon size using check primers for each locus. The *ompR234* allele was verified via sequencing of both strands of the amplicon.

Cell strains used in experiments in this study were created by transforming plasmids constructed above into MG1655 *PRO* Δ*csgA ompR234*, and are described in Supplementary Table 3.

### Culture conditions

Seed cultures were inoculated from frozen glycerol stock and grown in LB-Miller medium using appropriate antibiotics at the following concentrations: carbenicillin at 50 μg/ml, chloramphenicol at 25 μg/ml. Seed cultures were grown for 12 h at 37C in 14-ml culture tubes (Falcon), with shaking at 300 rpm. Experimental cultures were grown in M63 minimal medium (Amresco) supplemented with 1 mM MgSO_4_ and with 0.2% w/v glucose or 0.2% w/v glycerol. All liquid experimental cultures were grown in 24-well plate wells at 30C with no shaking. For inducing conditions, inducers used were anhydrotetracycline (Sigma) at concentrations of 1-250 ng/ml and N-(β-ketocaproyl)-L-homoserine lactone (Sigma) at concentrations of 1-1000nM. All experiments reporting standard error of the mean (s.e.m.) error bars were performed in triplicate.

### Transmission electron microscopy

For transmission electron microscopy (TEM), a 20 µl droplet of sample was placed on parafilm (Pechiney), and a 200-mesh formvar/carbon coated nickel TEM grid (Electron Microscopy Sciences) was placed with coated side face-down on the droplet for 30-60s. The grid was then rinsed with ddH_2_O by placing the grid face-down in a 30µl droplet of ddH_2_O and wicking off on filter paper (Whatman), and placed face-down for 15-30s on a droplet of 2% uranyl acetate (Electron Microscopy Sciences) filtered through 0.22 µm syringe filter (Whatman). Excess uranyl acetate was wicked off and the grid was allowed to air dry. Images were obtained on a FEI Tecnai Spirit transmission electron microscope operated at 80 kV accelerating voltage.

### Scanning electron microscopy

For scanning electron microscopy (SEM), biofilm samples on coverslips were coated with carbon sputtered to ∼10 nm with a Desk II Sputter Coater (Denton Vacuum). The samples were then imaged with a JEOL JSM-6010LA scanning electron microscope operated at 10 kV accelerating voltage. Images were obtained in secondary electron imaging (SEI) mode, and elemental mapping was performed with energy dispersive X-ray spectroscopy (EDS).

### AuNiNTA particle labeling assay

For nickel nitrilotriacetic acid gold nanoparticle (AuNiNTA particle) labelling of histidine tags displayed on CsgA, 200-mesh formvar/carbon coated nickel TEM grids (Electron Microscopy Sciences) were placed with coated side face-down on a 20µl droplet of sample on parafilm for 2 minutes. The side of the TEM grid with sample was rinsed with a 30 µl droplet of ddH_2_O, then with selective binding buffer (1X PBS with 0.487M NaCl, 80 mM imidazole, and 0.2v/v% Tween20), and placed face-down in a 60 µl droplet of selective binding buffer with 10nM 5 nm AuNiNTA particles (Nanoprobes). The TEM grid and droplet on parafilm was covered with a petri dish to minimize evaporation and allowed to incubate for 90 minutes. The grid was then washed 5 times with selective binding buffer without AuNiNTA particles, then twice with 1X PBS and twice with ddH_2_O. The thoroughly washed grid was placed face-down on a droplet of filtered 2% uranyl acetate for 15-30s to negative stain the sample. Excess uranyl acetate was wicked off with filter paper and grid allowed to air dry. Images were obtained on a FEI Tecnai Spirit transmission electron microscope operated at 80 kV accelerating voltage.

### Strain verification experiments (Supplementary Fig. 1, Fig. 1a)

For strain verification experiments, all cultures had an initial seeding concentration of 5X10^7^ cells/ml and were grown in glucose-supplemented M63. For experiments to verify that the fibrils seen on TEM were self-assembled from CsgA, MG1655 *ompR234* was grown for 24 h, aTc-Receiver/CsgA was grown for 14 h with 62.5 ng/ml aTc, and aTc-Receiver/CsgA-His was grown for 14 h with 62.5 ng/ml aTc. Biofilms resulting from aTc-Receiver/CsgA and aTc-Receiver/CsgA-His cultures were resuspended in 1XPBS and the AuNiNTA particle labelling assay (above) was performed to characterize curli fibrils. For experiments to verify that insertion of histidine tags into CsgA still allows curli fibril production, aTc-Receiver/CsgA was grown for 14 h with 250 ng/ml aTc, aTc-Receiver/CsgA-His was grown for 14 h with 250 ng/ml aTc, AHL-Receiver/CsgA was grown for 14 h with 1000nM AHL, and AHL-Receiver/CsgA-His was grown for 14 h with 1000nM AHL. The resulting biofilms were resuspended in 1XPBS and TEM imaging was performed.

Experiments were performed to verify that the MG1655 *PRO* Δ*csgA ompR234* host strain does not produce curli fibrils, that AHL-Receiver/CsgA-His only produces curli fibrils when induced by AHL, and that aTc-Receiver/CsgA-His only produces curli fibrils when induced by aTc. MG1655 *PRO* Δ*csgA ompR234* and MG1655 *ompR234* were grown for 48 h. AHL-Receiver/CsgA-His was grown for 48 h with no inducer, 1000nM AHL, or 250 ng/ml aTc. Similarly, aTc-Receiver/CsgA-His was grown for 48 h with no inducer, 1000nM AHL, or 250 ng/ml aTc. The resulting biofilms were resuspended in 1XPBS and TEM imaging was performed. ImageJ (NIH) was used to threshold images and calculate area covered by curli fibrils and area covered by cells. The ratio of areas was used to quantify curli fibril production.

### Crystal violet biofilm visualization (Supplementary Fig. 2)

Experiments were performed to verify that aTc-Receiver/CsgA-His formed thick adherent biofilms only when induced by aTc. Thermanox coverslips (Nunc) were placed at the bottom of 24-well plate wells in which aTc-Receiver/CsgA-His at an initial seeding density of 5X10^7^ cells/ml was grown for 24 h in glucose-supplemented M63 with no aTc or with 250 ng/ml aTc. Thermanox coverslips were immersed in 1% aqueous Crystal Violet (Sigma) for 20 minutes, after which the coverslips were washed by immersing in ddH_2_O repeatedly until no more visible dye was washed off. The Thermanox coverslips were allowed to air dry and digital photographs were taken.

### Conductive biofilm demonstration (Fig. 1b–d, Supplementary Fig. 3)

To create interdigitated electrodes (IDEs) for measurement of biofilm conductance, custom shadowing masks (Tech-Etch) with holes of the appropriate dimensions (Supplementary Fig. 4) were placed over Thermanox coverslips. Gold was sputtered to a thickness of ∼200 nm with a Desk II Sputter Coater (Denton Vacuum) and the resulting IDEs were tested with a Keithley 4200 picoammeter with two-point probe to confirm the absence of short circuits.

IDEs were placed at the bottom of 24-well plate wells and covered with glucose-supplemented M63 media with 250 ng/ml anhydrotetracycline (aTc), aTc-Receiver/CsgA-His cells at 1×10^8^ cells/ml, and 100nM AuNiNTA particles. The cultures were incubated at 30C with no shaking for 24 hours. For control experiments, either aTc was excluded, AHL-Receiver/CsgA-His cells were used in place of aTc-Receiver/CsgA-His cells, or AuNiNTA particles were excluded.

To obtain samples of the biofilm for TEM, a small amount of biofilm was scraped from the IDE substrate and resuspended in 1XPBS. IDEs were washed by repeatedly immersing in ddH_2_O, laid on a flat surface, and allowed to air dry for three days. A Keithley 4200 picoammeter with two-point probe was used to carry out a voltage sweep; a voltage difference was applied across the two probes and the current measured. Resulting I-V curves were fitted to simple linear regression lines and conductance of the biofilm was obtained from the slope. SEM imaging was then performed to characterize intact biofilm; SEM imaging was performed after conductance measurements because the samples were coated with a conductive carbon layer to facilitate imaging.

### Patterning experiments (Fig. 2–5, Supplementary Fig. 5–6)

In all cases, cultures were grown at 30C with no shaking. Also in all cases, after patterning experiment culture, the resulting biofilm was resuspended in 1X PBS and the AuNiNTA particle labelling assay above and TEM imaging were performed to characterize the patterned amyloid fibrils. ImageJ (NIH) was used to measure the length of unlabeled and AuNiNTA labeled fibril segments.

For tunable patterning based on changing induction time, AHL-Receiver/CsgA at an initial seeding density of 5 × 10^6^ cells/ml and aTc-Receiver/CsgA-His at an initial seeding density of 5X10^5^ cells/ml were co-cultured in 24-well plate wells with a 13 mm glass coverslip (Ted Pella) placed at the bottom. Cells were cultured in glycerol-supplemented M63 media. The cells were first co-cultured in the presence of 50nM N-(β-ketocaproyl)-L-homoserine lactone (AHL) inducer for 18-48 h. The media was then removed and the resulting biofilm incubated with no-inducer media for 6 h. The no-inducer media was then removed and replaced with media with 50 ng/ml aTc inducer. The biofilm was resuspended and the culture incubated for 16h. The resulting biofilm was resuspended and the AuNiNTA particle labelling assay (above) was performed to characterize curli fibrils. For the control experiment to produce fibrils when CsgA and CsgA-His were secreted slowly and simultaneously without temporal separation, AHL-Receiver/CsgA and aTc-Receiver/CsgA-His were co-cultured for 24 h in glycerol-supplemented M63 media with induction by 50nM AHL and 50 ng/ml aTc simultaneously.

For tunable patterning based on varying inducer concentration, AHL-Receiver/CsgA and aTc-Receiver/CsgA-His, each at an initial seeding density of 5X10^7^ cells/ml, were co-cultured in glucose-supplemented M63 media with AHL inducer at 0-1000nM and aTc inducer at 0–250 ng/ml, for 18 h.

For production of a dynamic material whose composition changes with time, a cell-cell communication system consisting of AHL-Sender+aTc-Receiver/CsgA and AHL-Receiver/CsgA-His was used. The two cell strains were co-cultured with AHL-Sender+aTc-Receiver/CsgA at an initial seeding density of 5X10^6^ cells/ml and AHL-Receiver/CsgA-His at an initial seeding density of 5X10^7^, 5X10^6^, or 5X10^5^ cells/ml. The cells were co-cultured in glucose-supplemented M63 media with 50 ng/ml aTc for 4-36 h. The resulting biofilm was resuspended and the AuNiNTA particle labelling assay was performed to characterize curli fibrils. For controls, AHL-Sender+aTc-Receiver/CsgA and AHL-Receiver/CsgA-His were grown separately. In each case, the cells were grown for 16 h with 50 ng/ml aTc induction, with an initial seeding density of 5X10^6^ cells/ml.

For patterning at two different length scales by combining genetic regulation of subunit expression with spatial inducer gradients, inducer-responsive cells were grown in solid phase on agar with opposing AHL and aTc inducer gradients. Inducer gradient agar plates were prepared by a two-step process^43^. A 100mmX100 mm square petri dish (Ted Pella) was elevated at one end by 1 cm. It was first filled by 20 ml of 1.5% agar glucose-supplemented M63 with 50nM AHL and was allowed to harden into an agar wedge. The plate was then laid flat and filled with 20 ml of 1.5% agar glucose-supplemented M63 with 50 ng/ml aTc. The plates were left for 12 h to allow diffusion of inducers to set up a concentration gradient. The surface of the agar was then overlaid with 5 ml of 0.7% agar glucose-supplemented M63 (top-agar) embedded with four cell strains–AHL-Receiver/CsgA, aTc-Receiver/CsgA-His, AHL-Receiver/GFP, and aTc-Receiver/mCherry – each at a cell density of 3.33X10^8^ cells/ml. The top-agar with embedded cells was allowed to harden, and the plate was incubated at 30C for 40 h. Fluorescence was imaged with a ChemiDoc MP imaging system (BioRad) and fluorescence intensity values extracted with ImageJ (NIH). Top agar was sampled and resuspended in 1XPBS, and the AuNiNTA particle labelling assay was performed to characterize curli fibrils. For the control condition with no inducer gradients, the same procedure was carried out but the petri dish was not elevated at one end to create agar wedges.

For patterning by protein-level engineering AHL-Receiver/8XCsgA-His, at an initial seeding density of 5X10^6^ cells/ml, was cultured in glucose-supplemented M63 with 10nM AHL inducer for 28 h. The resulting biofilm was resuspended and the AuNiNTA particle labelling assay was performed to characterize curli fibrils. For the control experiment, aTc-Receiver/CsgA-His at an initial seeding density of 1.5X10^6^ cells/ml was cultured in glucose-supplemented M63 with 10 ng/ml aTc inducer for 28 h.

For patterning at two different length scales by combining genetic regulation of subunit expression with subunit-level protein engineering, AHL-Receiver/8XCsgA-His at an initial seeding density of 5X10^7^ cells/ml and aTc-Receiver/CsgA-His at an initial seeding density of 5X10^5^ cells/ml were first co-cultured in glucose-supplemented M63 media with 4nM AHL inducer for 36 h. The media was then removed and the resulting biofilm incubated with no-inducer glucose-supplemented M63 media for 6 h. The no-inducer media was then removed and replaced with glucose-supplemented M63 media with 1 ng/ml aTc inducer. The biofilm was resuspended and the culture incubated for 18 h.

## Acknowledgements

We thank J.J. Collins (Biomedical Engineering, Boston University) for donating riboregulator plasmids, R. Weiss (Electrical Engineering and Computer Science, MIT) for the gift of a LuxI plasmid, and C. Dorel (Biosciences Department, INSA Lyon) for the gift of *E. coli* MG1655 OmpR234. We thank Z. Deng, B. Zakeri, C. Zhong, and P. Siuti from the Lu lab, and S. Keating from the lab of Neri Oxman (Media Lab, MIT) for helpful discussions. This work was supported by the Office of Naval Research and Army Research Office. This work was also supported in part by the MRSEC Program of the National Science Foundation under award number DMR-0819762. A.Y.C. acknowledges graduate research support from the Hertz Foundation, the Department of Defense, and NIH Medical Scientist Training Program grant T32GM007753. T.K.L. acknowledges support from the Presidential Early Career Award for Scientists and Engineers and the NIH New Innovator Award (1DP2OD008435).

## Author contributions

T.K.L. and A.Y.C. conceived the experiments. A.Y.C., U.O.S., M.Y.L., and R.J.C. performed the experiments, A.Y.C. and T.K.L. analysed the data, discussed results, and wrote the manuscript.

## Additional information

Correspondence and requests for materials should be addressed to T.K.L.

